# CM/Pf contribution to mesocircuit dynamics underlying awareness in non-human primates

**DOI:** 10.64898/2026.07.30.741726

**Authors:** Julie Maulavé, Florent Gobert, Justine Debatisse, Simon Clavagnier, Lisa Gauthier, Benjamin Pasquereau, Inés Mérida, Nicolas Costes, Luc Zimmer, Jacques Luauté, Maude Beaudoin-Gobert, Léon Tremblay

## Abstract

**Background:** In acute disorders of consciousness following severe brain injury, the heterogeneity and multiplicity of brain lesions make it difficult to infer a direct causal link between lesion patterns and impaired consciousness. As a result, neuroimaging analyses performed in intensive care settings can inform theories of consciousness mainly through correlational, rather than causal, evidence. The mesocircuit hypothesis proposes that interactions between the thalamus, basal ganglia, and cortex support the neural mechanisms underlying awareness. Among its key components, the intralaminar thalamic nuclei are thought to play a central role, yet their respective contributions to mesocircuit dynamics remain poorly understood.

**Methods:** In two macaque monkeys, we combined causal perturbations of three thalamic nuclei (the centromedian-parafascicular complex (CM/Pf), central lateral nucleus (CL), and mediodorsal nucleus (MD)) with a cue-guided behavioural task requiring the processing of multisensory information. Electrical microstimulation was first used to compare the behavioural consequences of perturbing each target. Reversible pharmacological inactivation using muscimol was subsequently performed in the CM/Pf and CL. Finally, the large-scale metabolic effects of CM/Pf inactivation (muscimol) and disinhibition (bicuculline) were assessed using [^18^F]FDG PET imaging.

**Results:** Perturbation of the intralaminar nuclei disrupted behavioural performance, with the strongest and most reproducible effects observed following CM/Pf manipulation. CM/Pf inactivation reduced correct responses and increased omissions and wrong responses, particularly under conditions requiring greater behavioural engagement. Importantly, these deficits occurred without evidence of major wakefulness impairment, as animals remained responsive throughout testing and showed no substantial increase in eye-closure probability. In contrast, CL perturbation produced weaker behavioural effects, suggesting distinct functional contributions of the two intralaminar nuclei. [^18^F]FDG PET imaging revealed that CM/Pf disinhibition increased glucose metabolism across thalamic, striatal, pallidal, and frontal cortical territories, whereas CM/Pf inactivation induced a broadly reciprocal pattern of metabolic pattern. These findings identify the CM/Pf as a major modulator of both thalamo-striatal and thalamo-cortical components of the mesocircuit.

**Conclusions:** Our results show that a unilateral intralaminar nucleus perturbation is not sufficient to affect wakefulness, suggesting that this function is mediated by a bilateral network at the thalamus level within the mesocircuit. The CM/Pf emerged as the intralaminar nucleus exerting the strongest influence on goal directed behaviours requiring the mesocircuit dynamics. Our findings also support the existence of partially distinct intralaminar pathways contributing differentially to global state regulation of awareness-related functions, and identify the CM/Pf as a key thalamo-striatal hub within the mesocircuit.

## Introduction

Acute brain injury in patients admitted in intensive care, consciousness may be transiently abolished by damage to strategic brain regions, resulting in severe disorders of consciousness (DoC). Such lesions classically involve either focal injury to single midline structures^1,2^ (possibly lateralised as the left-sided coma-causing-lesion^2^ of the brainstem) or bilateral networks, particularly within the diencephalon, as in bilateral thalamic infarction due to artery of Percheron occlusion.^3^ Alternatively, severe DoC may arise from diffuse telencephalic dysfunction, including diffuse white matter injury disrupting long-range cortico-cortical connectivity or widespread cortical injury.^4^ Predicting recovery and long-term functional outcome in DoC patients remains a major medical, ethical and social concern. Yet, identifying reliable biomarkers of awakening requires a deeper understanding of the neural mechanisms underlying consciousness.

In clinical practice, consciousness is often defined as a two-dimensional concept: arousal (i.e. the physiological level of wakefulness) and awareness (i.e. the content of consciousness, e.g. self- and environment perception, memory processes, communication).^5^ However, this simplified view is primarily operational and reflects only partially the richness of the semantic discussion surrounding contemporary theories of consciousness.

Contemporary theories such as the Global Neuronal Workspace (GNW) model propose that conscious access depends on the late, large-scale amplification and broadcasting of information across distributed fronto-parietal networks.^6,7^ In this theoretical view, consciousness is primarily defined by the global availability of specific contents for reportability and goal-directed behaviour, rather than by a unitary level of consciousness. This access-based account can be articulated with multidimensional components of consciousness, which distinguish global background conditions (such as wakefulness and large-scale cortical integration) from more selective impairments affecting access to specific contents or cognitive dimensions.^8^ Such a distinction is particularly relevant in DoC, where preserved behavioural wakefulness may coexist with impaired access to task-relevant information, corresponding to global/local states.^8^ It suggests that unilateral thalamic lesions may alter awareness-related processing without producing a global suppression of arousal. This conceptual framework therefore raises the question of how these distinct dimensions of consciousness are implemented by the underlying neural circuitry.

Cortico-striato-pallidal loops, traditionally associated with motor and cognitive functions, are now recognized as components of broader circuits that include outputs to thalamocortical systems (central thalamus and intralaminar nuclei) under brainstem modulation. These interconnected pathways form the so-called “mesocircuit”,^9^ proposed as a key substrate for widespread cortical arousal and the regulation of conscious processes. In the mesocircuit framework, consciousness is hypothesized to emerge from dynamic interactions between cortical and subcortical structures, supporting both arousal and conscious access. However, the respective contribution of its distinct components to wakefulness and awareness remains incompletely understood. The mesocircuit model^10^ provides an integrative framework for this architecture, linking thalamo-cortical loops with ascending arousal systems. In this model, the central thalamus occupies a pivotal position between ascending arousal systems supporting wakefulness and thalamo-cortical dynamics associated with awareness.^11^ Through this dual embedding, it may contribute not only to the regulation of global brain states, but also to the modulation of distributed cortical processes related to awareness. Thus, numerous studies have highlighted the key role of the striato-pallido-thalamic network in DoC.^10–13^ More specifically, pallidal and nigral GABAergic neurons project to the thalamus, notably to the thalamic reticular nucleus, where they play a crucial role in modulating thalamo-cortical circuits by exerting tonic and phasic inhibition on thalamic relay neurons, thereby influencing sensory processing, attention, emotion, and states of consciousness.

In light of the mesocircuit hypothesis,^10^ these observations raise the question of whether the thalamus modulates large-scale cortical information broadcasting. Whether these functions are uniformly supported by the thalamus or instead differentially mediated by distinct thalamic nuclei remains unknown. This view is supported by the largest multimodal meta-analysis of chronic DoC to date, combining structural, metabolic and functional imaging.^14^ The thalamus appears at the crossroads of brainstem arousal systems, basal ganglia loops, and widespread cortical territories, although this pivotal role is not unique, as previously illustrated by lesion network mapping studies of extra-thalamic sub-cortical areas in vascular coma.^15^

Within the thalamus, distinct associative and intralaminar nuclei may differentially contribute to the regulation of consciousness by acting either as relays of global states or as modulators of local conscious processing. The intralaminar thalamic nuclei, in particular the central lateral nucleus (CL) and the centromedian-parafascicular complex (CM/Pf), occupy a central position within the anterior forebrain mesocircuit, receiving ascending arousal-related inputs from the brainstem and basal forebrain and influencing global cortical excitability, behavioural responsiveness, wakefulness fluctuations, and the capacity of cortical networks to enter states compatible with conscious access. Accordingly, these central intralaminar nuclei would not necessarily generate specific local contents of consciousness, but would instead modulate the global state conditions under which such contents can emerge. By contrast, the mediodorsal thalamic nucleus (MD) is not an intralaminar nucleus, but a higher-order associative thalamic nucleus with dense reciprocal connections to the medial, orbitofrontal, dorsolateral prefrontal and anterior cingulate cortices, as well as subcortical limbic and fronto-striatal circuits. Rather than acting primarily as a relay of wakefulness, MD may contribute to the maintenance of awareness-related local states. This distinction is supported by recent lesion-mapping data in chronic DoC^14^ showing that thalamic compromise predominantly involves central thalamic regions, with a particularly high prevalence of MD involvement, observed in approximately 90% of cases, followed by CL in 38% and CM/Pf in 33%. These findings raise the possibility that different thalamic nodes may contribute to impaired consciousness through partially dissociable mechanisms: MD damage may reflect disruption of associative thalamo-prefrontal modulation of awareness-related local processing, whereas CL and CM/Pf damage may impair the global arousal and activation constraints required for such local processing to become functionally effective.

Although numerous neuroimaging studies have explored neuronal correlates of consciousness in DoC, correlational approaches linking heterogeneous lesion patterns to clinical status remain insufficient to establish causal relationships. To move beyond correlational inference, experimental models allowing controlled, reversible perturbations of candidate structures are required. Non-human primate (NHP) models provide such an experimental framework through causal neuromodulation. Previous studies have shown that stimulation of CL pathways can modulate attentional processes and executive control.^16^ Importantly, stimulation of central thalamic pathways has also been reported to restore wakefulness in anesthetized NHPs, accompanied by increased cortical metabolic and functional activity in the prefrontal, parietal, and cingulate regions.^17,18^ These findings further suggest that central thalamic modulation may influence both wakefulness and reversible states of consciousness. By contrast, the contribution of the CM/Pf has received considerably less attention despite its strategic position within the mesocircuit.

Previous studies have shown that the CM/Pf inactivation selectively alters attentional orienting while preserving overall wakefulness and basic sensorimotor responsiveness in macaques.^19^ Although only a limited number of studies have directly manipulated the CM/Pf pharmacologically in non-human primates, these findings consistently suggest a role in attentional orienting and the processing of behaviourally relevant sensory information.^20,21^ CM/Pf differs from CL in its projection patterns and functional integration within cortico-striatal loops, suggesting that distinct intralaminar nuclei may contribute differentially to parallel components of the mesocircuit organization. In contrast to CL, which projects diffusely to frontal and parietal cortices, the CM/Pf complex provides dense excitatory inputs to sensorimotor (posterior putamen) and associativo-limbic territories of the striatum (caudate nucleus and ventral striatum). Unlike the projection of the CL, which acts more directly on diffuse thalamo-cortical networks, this anatomical position of the CM/Pf complex could allow it to modulate mesocircuit activity through a thalamo-striato-pallidal pathway influencing behaviourally relevant cortical network engagement. This organization raises the possibility that the mesocircuit relies on partially parallel pathways, with CL preferentially interacting with thalamo-cortical dynamics related to arousal, and CM/Pf modulating striato-thalamo-cortical circuits involved in awareness. Consistent with this hypothesis, a recent systematic review of experimental and clinical evidence further highlighted the growing interest in specific intralaminar nuclei, particularly the CM/Pf complex, while concluding that their respective causal contributions to consciousness remain insufficiently understood.^22^ In parallel, central thalamic DBS has been explored in selected patients with prolonged DoC, with reports of behavioural improvement in some cases.^23^ However, clinical outcomes remain variable,^24^ and the mechanisms underlying these effects are not fully understood.

The present study aims to characterize the impact of perturbations of subcortical structures in non-human primates (NHP), focusing on the intralaminar nuclei of the mesocircuit (CL and CM/Pf) and the MD to test and compare the respective contribution of these three thalamic sites to a goal-directed task. It is therefore expected that focal disruption of cortico-striato-pallidal loops could produce alterations in behavioural responses. But how should we interpret and, therefore, name these changes in performance? Because unilateral thalamic disruption is not expected to substantially alter wakefulness,^25,26^ changes in goal-directed behavioural performance are interpreted here, for operational purposes, as modulations of awareness-related processes. This terminology acknowledges that the same effects may also be described in attentional, executive or motivational terms, but places them within the pragmatic wakefulness/awareness distinction^27^ interpreted within the theoretically sound local/global states framework.^8^ Accordingly, we test two alternative, although not mutually exclusive, hypotheses regarding the role of thalamic nuclei in consciousness-related functions to determine whether behavioural impairment is better explained by disruption of global-state regulation or by disruption of awareness-related local states. The first hypothesis predicts a predominant contribution of the intralaminar nuclei, whereby task impairment would mainly arise from altered global-state regulation through CL and/or CM/Pf dysfunction. The second hypothesis predicts a predominant contribution of MD, whereby impairment of the attentional task would reflect disruption of a thalamic node directly modulating the local state of consciousness associated with awareness-related cortical integration, in line with its dense connections with prefrontal, limbic, default-mode and fronto-parietal associative systems.

In brief, after identifying the thalamic targets producing the strongest behavioural effects during electrical stimulation, we performed muscimol (GABA agonist) microinjections in the CL and CM/Pf to reversibly enhance GABAergic inhibition. We compared muscimol and bicuculline (GABA antagonist) injections in the CM/Pf to assess their opposing effects on cerebral metabolism using imaging under anaesthesia in both monkeys.

## Materials and methods

### Animals and ethical statement

Two male monkeys (*Macaca fascicularis,* 5-6 years old, 5-8 kg body weight) were used in this study, housed in pairs in their home cage. They had *ad libitum* access to food, and controlled access to water, with supplementation of fruits and vegetables. The light-dark cycle (lights on from 08:00 AM to 7:00 PM), temperature (25°C) and humidity (60%) were kept constant in the animal room. Animal care and housing were compliant with National Institutes of Health guidelines (1996), the European Communities Council Directive 2010/63/UE (2010), and the French National Committee (87/848) recommendations, following the 3Rs rule (Reduction, Refinement, and Reuse or Rehabilitation). The procedure was approved by the French National Ethics Committee (#21774_2020011711591103). All experimental procedures were designed and conducted in accordance with the ARRIVE 2.0 guidelines.

### Surgery procedure and electrophysiological mapping for intra-thalamic stimulation and injection

A recording chamber and head-fixation system were implanted to enable stable head positioning and access to thalamic targets during experimental sessions. This implantation procedure has already been described in previous studies.^28,29^ Anatomical T1-weighted MRI scans (3T; CERMEP, Bron, France) were performed before surgery to stereotaxically position the recording chamber above the region of interest namely the posterior thalamus located 8.4 mm posterior to the anterior commissure. Detailed stereotaxic coordinates, chamber positioning, penetration depths, and electrophysiological mapping procedures are provided in the Supplementary Material.

### Behavioural task

Monkeys were trained to perform a behavioural task on a touchscreen placed in front of the monkey at a distance of 30 cm (Fig. 1A). Animals performed a cue-guided left-hand target selection task on a touchscreen, in which reward delivery depended on selecting the target associated with the sensory cue. Three conditions were used depending on the nature of the cue: visual, auditory and somaesthetic (see Supplementary Material for detailed task sequence, error categorization, sensory cue parameters, block structure, and stimulation timing).

**Figure 1:**
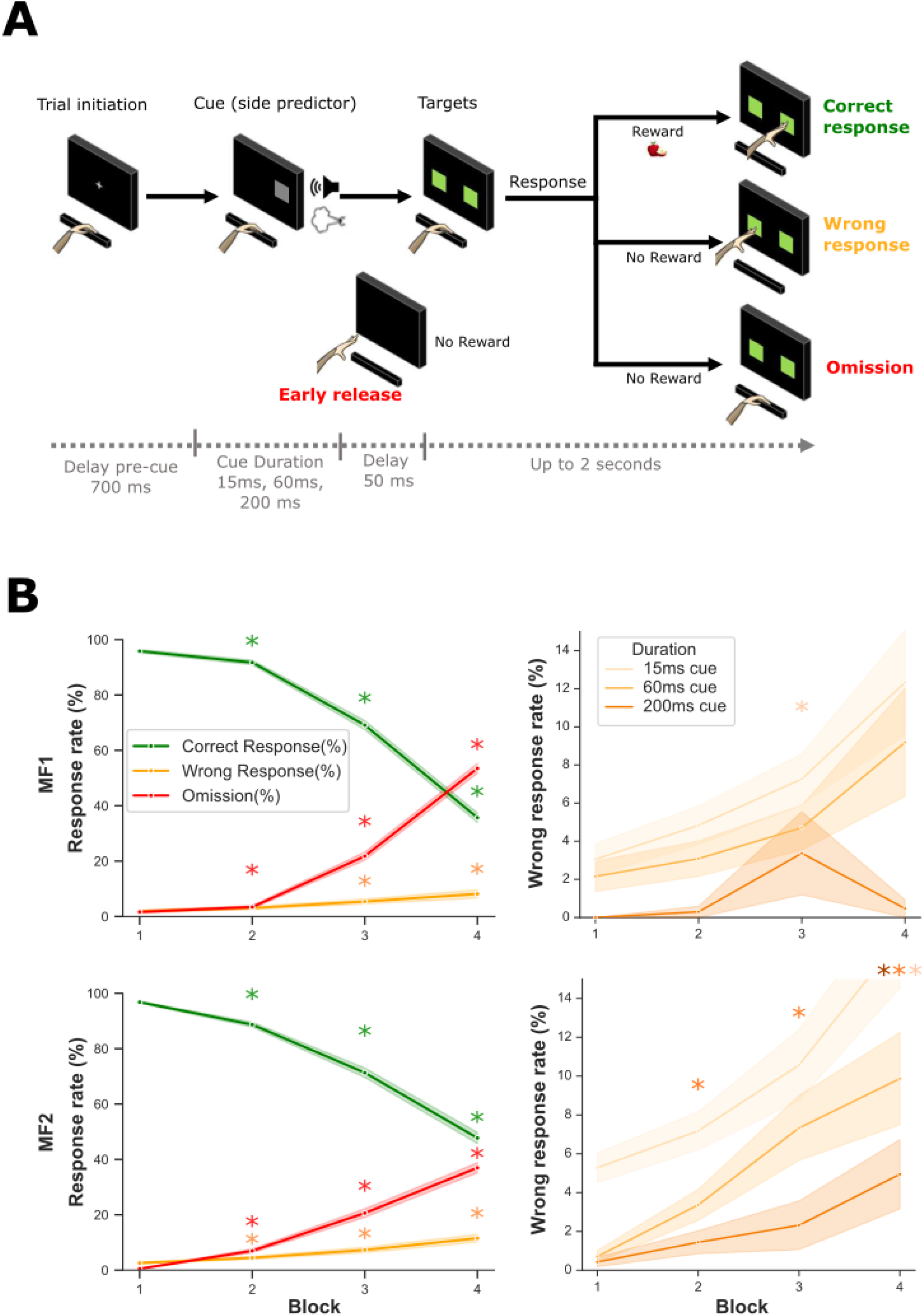
Behavioural task and performance during control sessions. **(A)** Schematic representation of the behavioural task. The monkey initiated each trial by placing its hand on a lever while facing a touchscreen. A lateralized cue was then presented on the left or right side of the screen. Depending on the condition, the cue consisted of a brief visual stimulus alone or combined with an auditory stimulus or an airpuff. Following cue presentation, two response targets appeared on the screen, and the animal was required to select the target corresponding to the cue location to obtain a liquid reward with the left hand, ipsilateral to the injection site to avoid any direct motor impairment in the response of the contralateral hemisphere (as the right handed response could have been more impaired by a direct alteration of the left-hemisphere motor loops). Incorrect target selection (wrong response), premature lever release (early release), non-target screen touch, or failure to respond within the allotted time (omission) were considered errors and were not rewarded. Each block consisted of 112 trials, and four consecutive blocks were performed during each session. **(B)** Behavioural performance during control sessions. Left: percentages of correct responses, wrong responses, and omissions across successive task blocks (mean ± SEM). Right: percentage of wrong responses for each animal as a function of cue duration (15, 60, or 200 ms) across blocks (mean ± SEM). Performance progressively deteriorated across successive blocks, with a decrease in correct responses (ANOVA, *F*(3,1818)=479.26, *P*<0.001) and increases in omission (*F*(3,1818)=379.74, *P*<0.001) and wrong responses (*F*(3,1812)=28.81, *P*<0.001). Post-hoc analyses showed significant differences between block 1 and blocks 2-4 for all three behavioural measures (all p<0.01). Shorter cue durations were associated with higher wrong response rates (*F*(2,1812)=42.50, *P*<0.001), and the effect of cue duration became more pronounced across successive blocks (bloc x cue duration interaction: *F*(6,1812)=2.44, *P*<0.05). These findings indicate that task performance is sensitive to both time-on-task and cue salience, providing a behavioural framework for evaluating the effects of thalamic manipulations.

Different parameters were measured, including reaction time, movement duration, the percentage of correct choices, the percentage of errors (including omission, early release and non-target touches) and wrong responses. The data are expressed as a percentage relative to the number of trials per block or per cue duration. A correct choice was defined as the selection of the correct target, previously identified by the cue. An early release corresponded to the premature removal of the hand from the lever. A wrong response corresponded to the selection of the incorrect target (i.e. the target opposite to that indicated by the predictive cue). An omission was defined as the absence of response. Detailed definitions of all error types are provided in the Supplementary Material.

Concerning the interpretation of these metrics: i) correct choices were analysed as an index of overall task performance; ii) wrong responses were examined to assess alterations in attentional engagement and behavioural selection, as they indicate that the animal was engaged in the task and able to perform the motor response but failed to select the correct target associated with the predictive stimulus; iii) omissions were analysed as an indicator of task engagement and represented the largest proportion of errors among the three error types.

### Eye-tracker recording

Eye tracker data were collected using a Spike 2 data acquisition system (Cambridge Electronic Design Ltd., CB, England). Eye state (open *vs* closed) was continuously monitored using an infrared eye-tracking system to estimate the probability of eye closure during task performance (see Supplementary Material for detailed signal processing and data analysis procedures).

### Microstimulation and muscimol microinjections during the behavioural task

Microstimulation was used to disrupt the three thalamic regions of interest (CM/Pf, CL and MD), and thus identify the targets for muscimol injections (Supplementary Fig. S1). Behavioural sessions were conducted either under control conditions or with unilateral thalamic microstimulation. During microstimulation sessions, microstimulation was delivered while animals performed the task. After the completion of the task, microstimulation sessions included an additional period alternating microstimulation and rest (see Supplementary Material). The stimulation sessions were conducted once daily, with one control session scheduled each week on a variable day. All stimulations were performed unilaterally, on the left side (see Supplementary Material for detailed stimulation site organization and coordinates).

The microinjections were performed according to the same protocol as in previous studies.^28,30^ All microinjections were performed unilaterally, on the left side. Muscimol (volume 1.5µL; concentration 1 µg/µL) was injected using a 30-gauge cannula tube connected to a 10 µL microsyringe (Hamilton) in the CM/Pf and CL nuclei, at the sites where microstimulation produced the strongest behavioural effects. Behavioural testing began ten minutes after the injection of muscimol. Detailed injection site organization and session scheduling are provided in the Supplementary Material.

### [^18^F]FDG PET imaging acquisition and analysis

PET-MRI acquisitions were performed at the imaging centre (CERMEP) under anaesthesia (alfaxan 10 mg/kg i.m. followed by propofol at a rate of 0,1 to 0,3 mg/kg/min i.v. 20 min later). Animals were ventilated and monitored. PET imaging was performed using a 3T Siemens Biograph mMR simultaneous PET-MRI scanner. Dynamic acquisition started with the IV injection of radiotracer fluorodeoxyglucose ([^18^F]FDG).

Each animal performed six PET-MRI scans: two controls, two during intrathalamic bicuculline microinjection and two during intrathalamic muscimol microinjections. Their heads were placed in a stereotaxic frame to perform microinjections. All microinjections were performed in the CM/Pf nuclei using stereotaxic coordinates determined during behavioural task. Bicuculline methiodide (volume 1 µL; concentration 15 µg/µL) and muscimol (volume 1.5 µL; concentration 1 µg/µL, identical to that used during the behavioural task) were injected 15 minutes before the injection of [^18^F]FDG.

Regional [^18^F]FDG standardized uptake value ratios (SUVR) were extracted from 92 regions of interest (ROIs) using an updated version of the *Macaca fascicularis* maximum probability atlas^31^ (Supplementary Fig. S2). Further details regarding image acquisition, preprocessing, and atlas construction are provided in the Supplementary Material.

### Statistical analysis

All statistical analyses were performed using Python, except for eye tracker analysis, which were performed using MATLAB. For control conditions, one-factor ANOVAs were performed (factor: block), separately for each behavioural parameter (correct trial rate, miss rate and wrong response rate), in order to assess performance changes across the different blocks of the session. One-factor ANOVAs were performed (factor: cue duration: 15, 60 or 200 ms), separately for each of the three behavioural parameters. A two-factor ANOVA (cue duration × block) was performed for the wrong response rate in order to evaluate the main effects and their interaction. A one-factor ANOVA (factor: cue type) was also performed on the wrong response rate.

For microstimulation, one-factor ANOVAs were performed (factor: stimulation territory *versus* control), separately for each of the three behavioural parameters. A three-factor ANOVA (animal × stimulation territory × block) was performed for each of the three behavioural parameters to assess the main effects and their interactions. For microinjection, one-factor ANOVAs were performed (factor: injection territory *versus* control), separately for each of the three behavioural parameters. A two-factor ANOVA (injection territory and control × cue duration) was performed for each of the three behavioural parameters. Unless otherwise specified, statistical analyses were performed on pooled data from both animals, whereas figures display individual data for each monkey.

Main and interaction effects were considered significant at *P*< 0.05. Within each ANOVA, post hoc analyses were conducted using two-sided Mann-Whitney-Wilcoxon tests. Data are represented as mean ± SEM.

## Results

### Control performance progressively declines with increasing attentional demand

During the control condition, the performances of the two animals significantly decreased during each daily session, across the successive task blocks. This was reflected in the ANOVA assessing the effect of block on correct trial rates (*F*(3,1818)=479.26; *P*<0,001), omission rates (*F*(3,1818)=379.74; *P*<0.001) and wrong responses rates (*F*(3,1812)=28.81; *P*<0.001) for both monkeys (Fig. 1B). Post hoc analysis revealed that correct trial rates were significantly lower in blocks 2 to 4 than in block 1 (all *P*<0.001). Omission rates were significantly increased in blocks 2 to 4 compared to block 1 (all *P*<0.001). Wrong responses rates were significantly increased in blocks 2 to 4 compared to block 1 (*P* min<0.001, *P* max<0.01; Fig. 1B).

Performances of both monkeys were also modulated by cue duration (Fig. 1B and Supplementary Fig. 3). There were no interactions between animal and cue duration (*F*(2,1818)=0.37, *P*=0.69) (Fig. 1B and Supplementary Fig. 1). Shorter cue durations were associated with a significantly higher percentage of wrong responses (*F*(2,1812)=42.50, *P*<0.001). Post hoc analysis revealed that wrong responses were significantly higher for 15 ms cues compared to 200 ms cues, for 15 ms cues compared to 60 ms and for 60ms cues compared to 200 ms (all *P*<0.001; Supplementary Fig. 3).

A significant two-way interaction was found between cues duration and block for the wrong responses (*F*(6,1812)=2.44, *P*<0.05). This effect appeared to be more pronounced in blocks 2 to 4 of the task than in block 1 (*P* min<0.001; *P* max>0.05; Fig. 1B). A significant main effect of cue type on wrong responses was observed (*F*(2,1812)=3.03, *P*<0.05). However, subsequent post hoc pairwise comparisons failed to identify significant differences between any specific cue types (all *P*>0.05) (Supplementary Fig. 3).

### Microstimulation identifies CM/Pf as the intralaminar nucleus with the strongest behavioural impact

To determine the behavioural impact of stimulating within the different thalamic nuclei, electrical microstimulation was performed within the CL, CM/Pf and MD nuclei, at six different sites for CL and MD, and four different sites for CM/PF per animal. An ANOVA revealed a significant main effect of stimulation territory for each behavioural parameter: correct choice rates (*F*(3,2425)=117.71; *P*<0.001), omission rates (*F*(3,2425)=94.04; *P*<0.001), and wrong responses rates (*F*(3,2425)=33.50; *P*<0.001).

Microstimulation of the CM/Pf resulted in a significant decrease in correct response rates compared to control (*P*<0.001 and *U*=96396.5; Fig. 2A). This was accompanied by a significant increase in omission rates (*P*<0.001 and *U*=219619; Fig. 2B) and wrong responses rate (*P*<0.001 and *U*=215405; Fig. 2C) compared to control. Microstimulation of the CL, significantly reduced correct choice rate compared to control (*P*<0.001 and *U*=337176.5; Fig. 2A), while increasing omission rate (*P*<0.001 and *U*=189037.5; Fig. 2B) and wrong responses rate (*P*<0.001 and *U*=201166.5; Fig. 2C). Microstimulation in the MD region led to a significant reduction in correct choice rate (*P*<0.001 and *U*=279133; Fig. 2A) and an increase in omission rate compared to control (*P*<0.001 and U=212364; Fig. 2B). However, the effects on wrong responses rates were not significant (*P*=0.186 and *U*= 241035.5; Fig. 2C).

**Figure 2:**
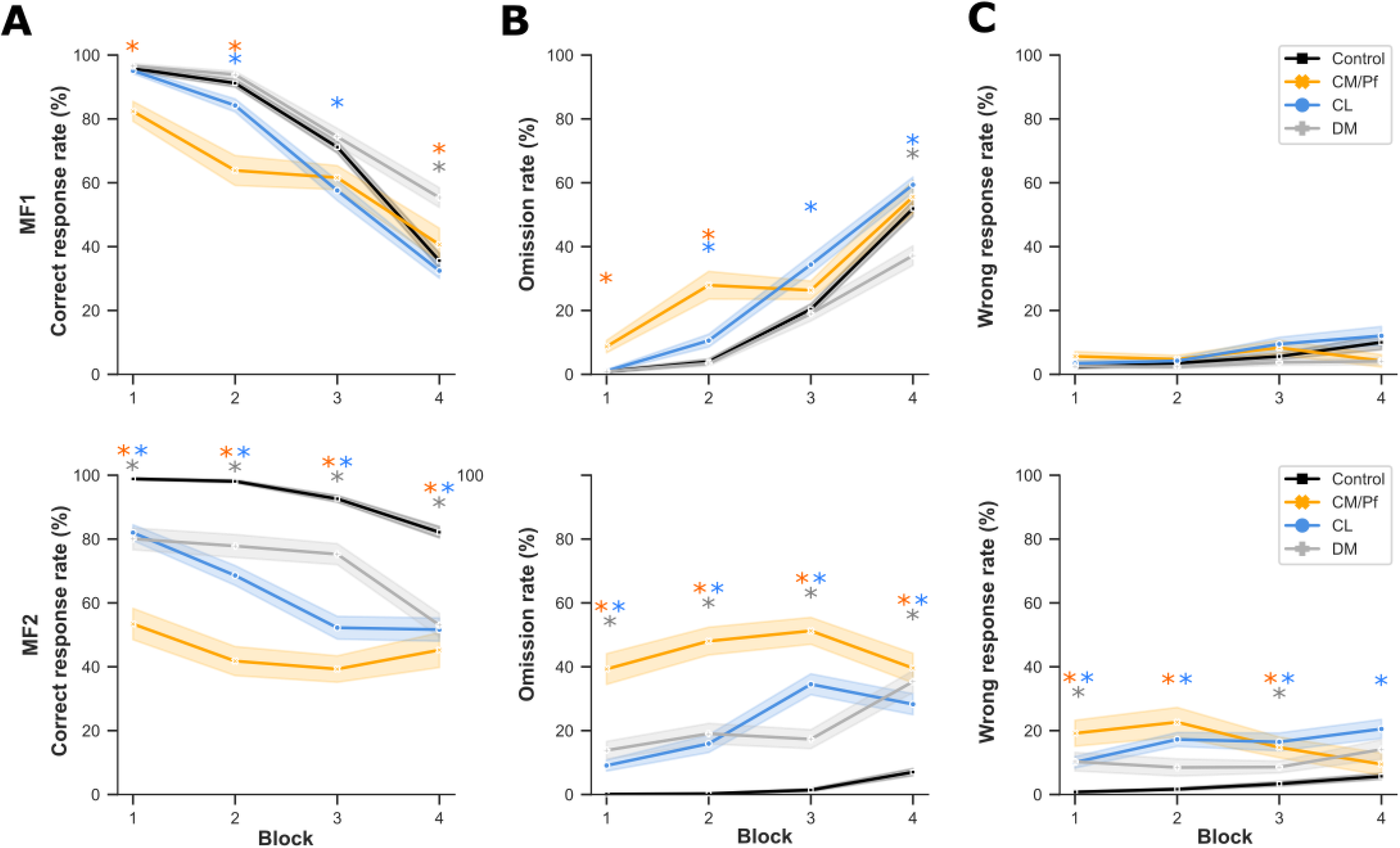
CM/Pf and CL microstimulations induce the strongest disruption of behavioural performances. **(A-C)** Behavioural performance during microstimulation of the centromedian-parafascicular complex (CM/Pf), central lateral nucleus (CL), or mediodorsal nucleus (MD). Data are shown separately for each animal across successive task blocks (mean ± SEM): **(A)** Percentage of correct choices, **(B)** Percentage of omissions, and **(C)** Percentage of wrong responses. A significant animal x stimulation territory x block interaction was observed for all behavioural measures (correct choices: *F*(21,225)=13.14, *P*<0.001; omissions: *F*(21,2425)=17.23, *P*<0.001; wrong responses: *F*(21,2425)=2.30, *P*<0.001), indicating that stimulation effects varied across animals and task progression. A significant main effect of stimulation territory was observed for correct choices (ANOVA, *F*(3,2425)=117.71, *P*<0.001), omissions (F(3,2425)=94.04, *P*<0.001), and wrong responses (*F*(3,2425)=33.50, *P*<0.001). Compared with control sessions, microstimulation of both the CM/Pf and CL significantly reduced correct choice rates and increased omission and wrong response rates (all *P*<0.001, two-side Mann-Whitney-Wilcoxon tests). MD stimulation produced weaker behavioural effects, increasing omissions and reducing correct choices without significantly affecting wrong response rates. Post hoc comparisons revealed that CM/Pf stimulation induced the strongest behavioural disruption, producing greater reductions in correct choices and greater increases in omissions than either CL or MD stimulation. CL stimulation also produced significantly greater behavioural impairment than MD stimulation. Overall, CM/Pf stimulation produced the most pronounced impairment in task performance, consistent with a critical contribution of this nucleus to behavioural engagement during demanding attentional processing.

A significant three-way interaction was found between animal, stimulation territory and block for each parameter (correct choice rate: *F*(21,225)=13.14; *P*<0.001; omission rate: (*F*(21,2425)= 17.23; *P*<0.001; wrong response rate: (*F*(21, 2425)=2.30; *P*<0.001), indicating that the stimulation effects varied between animals over time. Post hoc comparisons also showed that stimulation of the CM/Pf induced the strongest behavioural effect, compared to the CL and to the MD, by reducing correct choices rates (compared to the CL: *P*<0.05 and *U*=120970 and compared to the MD: *P*<0.001 and *U*=94919), increasing omission rates (compared to the CL: *P*<0.001 and *U*=150099 and compared to the MD: *P*<0.001 and *U*=160810) and increasing wrong response rates (compared to the MD: *P*<0.001 and *U*=144441.5 and compared to the CL: *P*=0.1066 and *U*=125949.5). CL stimulation also produced significantly greater behavioural effect than MD stimulation, reducing correct choices rates (*P*<0.001 and *U*=166783.5) and increasing omission rates (*P*<0.001 and *U*=230365) and wrong response rates (*P*<0.001 and *U*=243704).

Based on the stronger, consistent, and similar behavioural effects observed across monkeys following stimulation the CM/Pf and CL nuclei, in contrast to the weaker and more variable effects observed in the MD, subsequent pharmacological muscimol injections were performed in the CM/Pf and CL, whereas the MD was not further investigated.

### CM/Pf inactivation by microinjection preferentially impairs behavioural performance under high attentional demand

To further probe the causal effect of both intralaminar nuclei (CL and CM/PF), microinjections of muscimol, a GABA_A_ agonist, were used to reversibly enhance inhibitory tone and suppress the activity of these thalamic regions, at two different injection sites for the CM/Pf and three for the CL, with a total of eight injections per animal. An ANOVA revealed a significant main effect of injection territory for each behavioural parameters, correct choice rate (*F*(2,1794)=190.65; *P*<0.001), omission rate (*F*(2,1794)=85.51; *P*<0.001), and wrong response rate (*F*(2,1794)=24.37; *P*<0.001) (Fig. 3).

**Figure 3:**
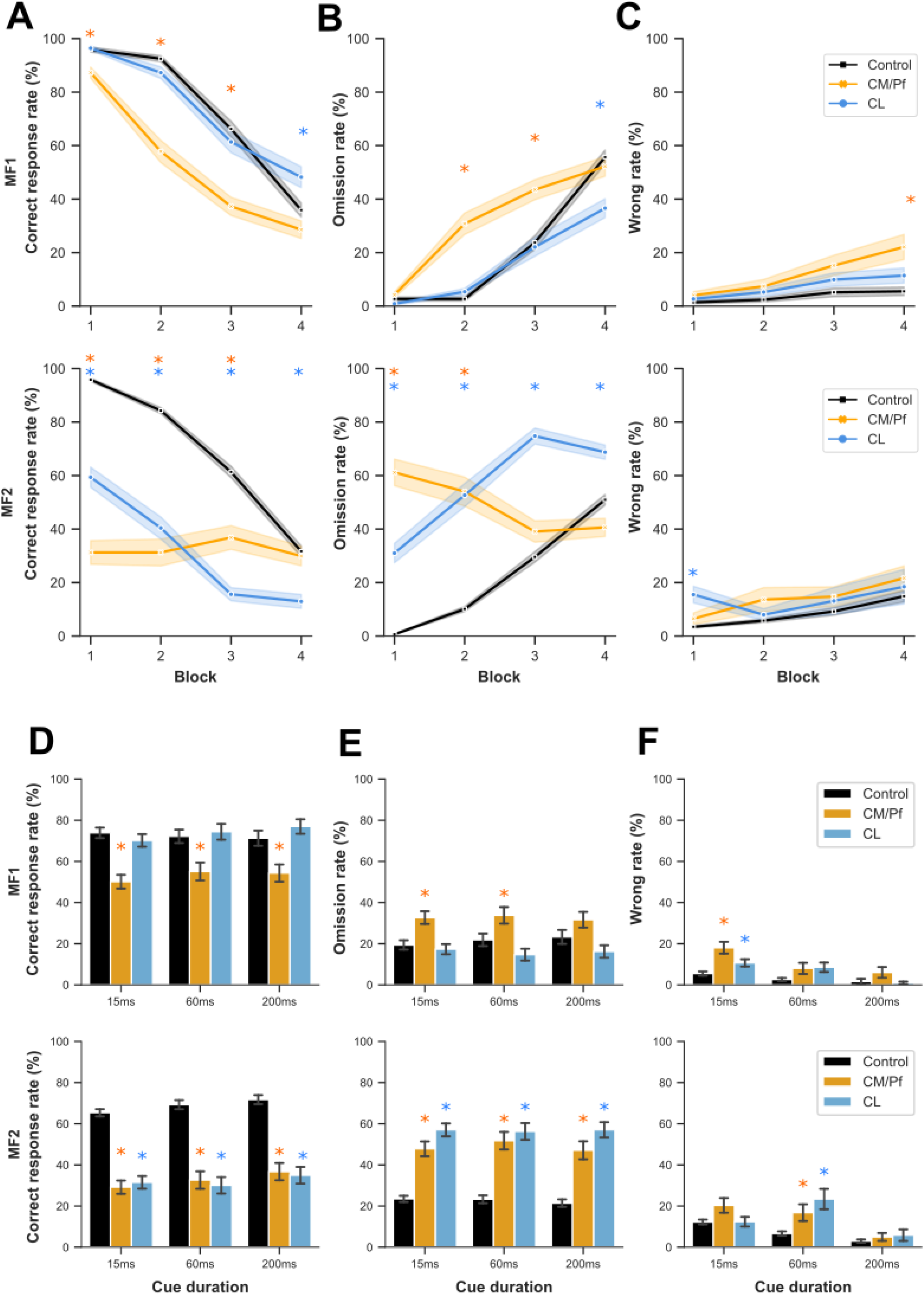
Reversible inactivation of CM/Pf and CL impairs behavioural engagement, with stronger effects following CM/Pf inactivation. **(A-C)** Behavioural performance following muscimol microinjections into the CM/Pf or CL, shows separately for each animal across successive task blocks (mean ± SEM): **(A)** Percentage of correct choices, **(B)** Percentage of omissions, and **(C)** Percentage of wrong responses. **(D-F)** Behavioural performance as a function of cue duration (15, 60, or 200 ms): **(D)** Percentage of correct choices, **(E)** Percentage of omissions, and **(F)** Percentage of wrong responses. A significant main effect of injection territory was observed for correct choices (*F*(2,1794)=190.65, *P*<0.001), omissions (*F*(2,1794)=85.51, *P*<0.001), and wrong responses (*F*(2,1794)=24.37, *P*<0.001). Compared with control sessions, muscimol injections into both CM/Pf and CL significantly reduced correct choice rates and increased omission rates (all *P*<0.001). Both manipulations also increased wrong response rates (CM/Pf: *P*<0.001; CL: *P*<0.01). A significant interaction between injection territory and cue duration was observed for wrong responses (*F*(4,1794)=5.40, *P*<0.001). CM/Pf inactivation preferentially increased wrong response rates at the shortest cue duration (15 ms; *P*<0.01 *versus* control), whereas longer cue durations were not significantly affected. Overall, behavioural impairments were most pronounced under conditions of increased attentional demand, indicating that CM/Pf perturbation disrupted behavioural access to task-relevant information rather than inducing a global reduction in responsiveness.

Muscimol microinjection into the CM/Pf resulted in a significant decrease in the correct choice rate compared to control at the group level (*P*<0.001 and *U*=122373.5; Fig. 3A). This was accompanied by a significant increase in omission rates (*P*<0.001 and *U*=269373.5; Fig. 3B) and wrong response rates (*P*<0.01 and *U*=226272; Fig. 3C) rate compared to control. CM/Pf inactivation resulted in significantly increased wrong response rates at the shortest cue duration compared to control (15ms cue: *P*<0.01 and *U*=42889.5; Fig. 3F). Longer cues durations (60ms cue and 200ms cue) did not significantly differ from control (respectively *P*=0.1276 and *U*=18444.5 and *P*=0.0902 and *U*=18024). These findings indicate that CM/Pf perturbation preferentially impaired behavioural processing under conditions requiring higher attentional engagement, rather than inducing a global reduction in wakefulness or responsiveness. Eye-tracker monitoring did not reveal major increases in eye-closure probability following CM/Pf inactivation compared to control conditions, arguing against a generalized reduction in wakefulness or increased drowsiness during task. These findings support the interpretation that behavioural alterations primarily reflected impaired attentional engagement and behavioural access to task-relevant information rather than global loss of wakefulness.

In comparison, microinjection in the CL significantly reduced correct choice rates compared to control (*P*<0.001 and *U*=161620; Fig. 3A), while increasing omission rates and wrong response rates (respectively *P*<0.001 and *U*=233942.5 and *P*<0.01 and *U*=216025; Fig. 3B-C). No significant effects of interaction between injection territory and cue duration were found for correct choice rates (*F*(4,1794)=0.359; *P*=0.838) and omission rates (*F*(4,1794)=0.347; *P*=0.386). A significant two-way interaction was found between injection territory and cue duration for wrong response rates only (*F*(4,1794)=5.40; *P*<0.001; Fig. 3D-F). CL inactivation resulted in significantly increased wrong response rates at 60 ms compared to control (*P*<0.001 and *U*=19178; Fig. 3F). Results were not significant at 15 ms and 200 ms compared to control (*P*=0.2765 and *P*=0.9945). Wrong response rates were significantly higher at 60 ms than at 200 ms (*P*<0.001 and *U*=4096.5). However, the comparison between 15 ms and 60 ms was not significant (*P*=0.9945). Overall, the effects of muscimol injections were most pronounced at shorter cue durations (between 15 and 60 ms depending on site injection), which were associated with higher wrong response rates.

### CM/Pf modulation engages widespread frontal, striatal and thalamic networks

Following local CM/Pf activation (by bicuculline microinjection), glucose metabolism was altered in several cortical and subcortical territories including the basal ganglia and the thalamus (Fig. 4). At the subcortical level, the strongest metabolic increase was observed at the injection site itself, within the CM/Pf, along with increases in several thalamic nuclei including the ventral anterior (VA), ventral lateral (VL), and ventral posterior medial (VPm) (Fig. 4A-B). Within the striatum, [¹⁸F]FDG uptake was elevated bilaterally in the ventral striatum, anterior caudate nucleus, anterior and posterior putamen, and external pallidum (GPe). At the cortical level, increased glucose metabolism was observed bilaterally in the medial prefrontal cortex (area 10), anterior cingulate cortex (areas 24 and 32), limbic cingulate cortex (area 25), medial and lateral orbitofrontal cortex (areas 13 and 12/45), and insula. Notably, contralateral increases were restricted to the limbic and orbitofrontal cortices, anterior cingulate cortex, anterior putamen, ventral striatum, and external pallidum, whereas ipsilateral changes were more widespread across frontal, striatal, and thalamic territories (Fig. 4B).

**Figure 4:**
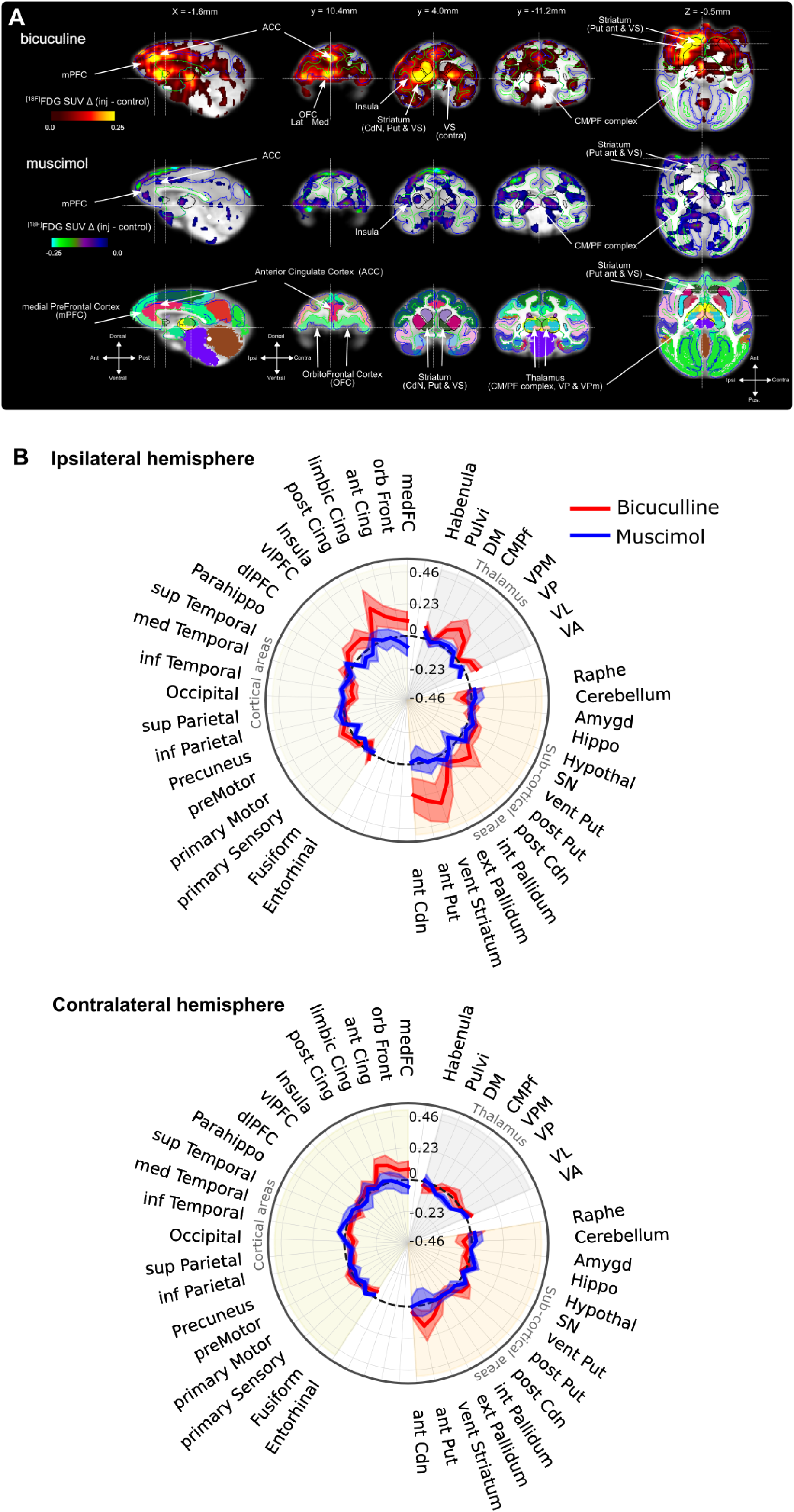
Opposing effects of CM/Pf disinhibition and inactivation on cortico-striato-thalamic metabolism. **(A)** Representative [^18^F]FDG-PET imaging co-registered to individual MRI and displayed in template space. Upper row: bicuculline injection into CM/Pf. Middle row: muscimol injection into the CM/Pf. Lower row: updated MaxProb Atlas^22^ used for anatomical localization (see Supplementary Fig. 3). **(B)** Regional standardized uptake value ratio (SUVR) differences relative to the control condition measured 75-90 min after injection. Values are shown separately for ipsilateral and contralateral hemispheres. Bicuculline-induced disinhibition of the CM/Pf increased glucose metabolism within the ipsilateral thalamus, bilateral basal ganglia, and ipsilateral frontal cortical regions, including the ventral striatum, caudate nucleus, putamen, pallidum, anterior cingulate cortex, orbitofrontal cortex, medial prefrontal cortex, and insula. In contrast, in muscimol-induced inactivation produced an overall reduction of glucose metabolism across largely overlapping cortico-striato-thalamic territories. These opposite metabolic effects identify the CM/Pf as a key modulator of widespread cortico-subcortical networks implicated in behavioural engagement and access to task-relevant information. *ant CdN: anterior Caudate Nucleus; ant Put: anterior Putamen; CM/Pf: Centromedian-Parafascicular; ext Pallidum: external Pallidum; Hypothal: Hypothalamus; MD: Mediodorsal; int Pallidum: internal Pallidum; post CdN: posterior Caudate Nucleus; post Put: posterior Putamen; Pulvi: Pulvinar; vent Str: ventral Striatum; VA: Ventral Anterior; VL: Ventral Lateral; VP: Ventral Posterior; VPM: Ventral Posterior medial; Raphe: Raphe nucleus. The orientation directions are illustrated by the arrows (D= dorsal; A= anterior, L= left)*.

Local CM/Pf inactivation (by muscimol microinjection) produced a broadly reciprocal pattern, reducing [¹⁸F]FDG uptake in several regions that had shown increased metabolism under bicuculline (Fig. 4A). Metabolic decreases were most pronounced ipsilaterally in the medial prefrontal cortex, dorsolateral prefrontal cortex (dlPFC), and insula, as well as bilaterally in the VL and ventral posterior (VP) thalamic nuclei and anterior putamen, and ipsilaterally in the posterior putamen (Fig. 4B).

## Discussion

The main objective of this study was to determine, in non-human primates, using complementary causal approaches (microstimulation and microinjection of GABAergic agents), which of the three thalamic regions proposed as key components of the mesocircuit modulation for local (MD) and global (CM/Pf and CL) states of consciousness most strongly modulates performance during a behavioural task probing multisensory experience integration, attention, motivation and behavioural responsiveness. Based on the mesocircuit framework presented in the introduction, we compared the respective contributions of the MD, CL and CM/Pf to determine whether these thalamic nuclei differentially modulate behavioural performance. Having identified the CM/Pf as the most behaviourally relevant thalamic target, we next investigated the large-scale metabolic consequences of its modulation using [¹⁸F]FDG PET imaging under propofol anaesthesia, focusing on the CM/Pf selected through the sequential behavioural screening (microstimulation followed by pharmacological inactivation).

Four principal findings emerged from this study: (1) combining intrathalamic microstimulation with a behavioural task using multisensory cues allowed the respective contribution of the three thalamic targets to be dissociated; (2) across both causal perturbation approaches, the CM/Pf complex consistently emerged as the intralaminar nucleus exerting the strongest influence on behavioural performance, whereas CL and MD produced weaker or less reproducible effects; (3) CM/Pf modulation engaged widespread frontal, striatal, thalamic and pallidal components of the mesocircuit, as revealed by the combination of pharmacological perturbations and [¹⁸F]FDG PET imaging and (4) combining behavioural and metabolic approaches provided complementary evidence supporting a prominent contribution of the CM/Pf within mesocircuit dynamics.

### Dissociating the functional roles of intralaminar thalamic nuclei

Our behavioural task allowed several parameters to be measured, including motivation (through movement duration, reaction time, omission rate and completion rate), impulsivity (through the analysis of early releases), and attentional processes, mainly assessed through the analysis of wrong response rates and dependent on cue durations.

During the baseline period, as the task progresses and motivation declines, animals show reduced attention, resulting in increased wrong responses and omissions, particularly for short-duration cues that require higher attentional engagement. These patterns indicate that the task was sensitive to fluctuations in behavioural state and processing demands, providing a suitable framework for assessing the consequences of thalamic perturbations (Fig. 1B).

Transient microstimulation of each thalamic target during the cue-guided task revealed that stimulation of the intralaminar nuclei (CL and CM/Pf) reduced task performance, as reflected by lower correct response rates and increased omission rates from the earliest task blocks of both animals. These effects were strongest following CM/Pf stimulation, identifying this nucleus as the intralaminar target exerting the greatest influence on task performance. In addition, CM/Pf stimulation produced the highest wrong response rates in MF2 from the earliest task blocks, consistent with a greater disruption of task performance, without altering basic sensorimotor capacities, as the animal remained able to generate behavioural responses. These findings supported the selection of the CM/Pf complex for the subsequent pharmacological inactivation using muscimol in the second part of the study. In contrast, MD stimulation produced weaker and less consistent behavioural effects across animals, supporting our decision not to pursue this nucleus in subsequent pharmacological experiments.

### Interpreting behavioural effects within the mesocircuit framework

Our experimental design was based on the assumption that unilateral modulation of thalamic nuclei involved in global states of consciousness would be insufficient to abolish wakefulness because of compensatory mechanisms within bilateral ascending arousal systems. Consistent with this assumption, the observed perturbations did not appear to interfere with global wakefulness, supporting a dissociation between preserved wakefulness and impaired higher-order conscious processing.

This interpretation is consistent with the experimental findings of Redinbaugh et al.,^17^ who showed that unilateral stimulation of the central lateral thalamus was sufficient to modulate arousal under experimental conditions, while clinical lesion studies indicate that sustained impairment of wakefulness is more commonly associated with bilateral paramedian or intralaminar thalamic lesions, particularly when extending to the rostral brainstem.^3^ Together, these observations support the view that unilateral perturbation of intralaminar thalamic nuclei may substantially alter higher-order behavioural performance while leaving global wakefulness largely preserved.

Because interpreting mesocircuit perturbations in terms of consciousness remains debated in preclinical models, including non-human primates,^27^ we explicitly define the conceptual framework adopted throughout this Discussion. A goal-directed behavioural task can legitimately be interpreted in terms of attentional, executive, and motivational dimensions, depending on the cognitive-science framework used. However, it can also be situated within the classical clinical distinction between wakefulness and awareness, despite the theoretical limitations of this dichotomy, as it provides an operationally useful framework for the present study. In this context, the proposed protocol was not expected to induce a substantial alteration of wakefulness (see above). Consequently, when consciousness-related modulations are observed without major wakefulness disruption, they may be more appropriately related to changes in the content of consciousness, a dimension that is often operationally encompassed by the term awareness in English-speaking clinical and cognitive frameworks. For this reason, modulations of cognitive task performance are here considered, for operational purposes, as modulations of awareness.

The theoretical interpretation of these findings depends on the thalamic target involved. Effects associated with mediodorsal thalamic modulation are interpreted as supporting a local-state hypothesis, in which MD influences the task-relevant conscious process through thalamo-prefrontal, limbic, default-mode and fronto-parietal associative networks. Conversely, effects associated with modulation of the intralaminar nuclei, namely the central lateral nucleus and the centromedian-parafascicular complex, are interpreted within a global-state hypothesis, in which CL or CM/Pf influence awareness-related performance indirectly by regulating arousal, attentional readiness and cortical excitability, rather than by directly generating the task-specific content of consciousness.

### CM/Pf perturbation selectively alters awareness-related behavioural performance

To further compare the respective contributions of the two intralaminar nuclei, we reversibly inactivated the CL and CM/Pf using muscimol during the cue-guided attentional task. This manipulation confirmed that CM/Pf perturbation produced stronger and more reproducible behavioural effects than CL perturbation. Unilateral inactivation of the CM/Pf complex selectively altered task performance without affecting basic sensorimotor abilities. The animals’ ability to respond with the ipsilateral hand relative to the injected hemisphere (left hand, ipsilateral to the injection site) was preserved, indicating that increased inhibitory tone did not impair basic sensorimotor capacities provided that the contralateral motor response was not required.

The effects observed following CM/Pf muscimol injections are consistent with an attentional deficit, as wrong choices occurred significantly more frequently. This increase indicates that animals were still able to detect the targets and execute the required movement but failed to appropriately use the sensory cue predicting the rewarded site. Importantly, these behavioural deficits occurred without evidence of major wakefulness impairment. Eye tracking further confirmed the absence of increased eye-closure probability during task performance, supporting the conceptual framework outlined above, whereby preserved wakefulness can coexist with impaired awareness-related performance.

Within the Global Neuronal Workspace (GNW) framework,^32^ this pattern is better interpreted as a selective impairment of conscious access to task-relevant contents, rather than as a disruption of wakefulness or of consciousness as a whole. Preserved arousal and behavioural responsiveness suggest that the global enabling conditions for consciousness were not abolished. However, the observed behavioural deficits may reflect a reduced probability that specific, behaviourally relevant representations are amplified, stabilized and made globally available within large-scale cortical networks. This interpretation remains compatible with Naccache’s access-consciousness account of the GNW,^33^ in which conscious processing depends on the all-or-none availability of information for reportability, although reportability itself was not directly assessed in the present study and behavioural responsiveness was used as its operational surrogate. In parallel, the terminology proposed by Bayne, Hohwy and Owen^8^ would describe these findings as affecting a specific dimension of conscious processing, rather than a unitary level of consciousness. In this framework, CM/Pf dysfunction would not be interpreted as directly generating a local state of consciousness, but rather as altering the thalamo-striatal and thalamo-cortical gating mechanisms that allow task-relevant information to gain access to the workspace and to be translated into consistent, goal-directed behavioural responses.

To our knowledge, only two studies have used muscimol to inhibit the CM/Pf complex in non-human primates, and both reported comparable behavioural findings. Minamimoto and colleagues reported a significant increase in reaction time for correctly cued targets presented contralateral to the injection site, indicating that the CM/Pf complex plays an essential role in directing attention toward contralateral external events.^19^ Similarly, Matsumoto and colleagues showed that muscimol injections altered monkeys’ licking behaviour, suggesting that CM/Pf inactivation disrupts the attentional processing of behaviourally relevant stimuli.^20^ Additional NHP studies support these findings, showing that lesions within the central thalamus can induce persistent disturbances in arousal regulation and cognitive deficits.^34^

The CM/Pf is the principal thalamic output to the striatum. Through this major thalamostriatal projection, it can regulate the activity of the striatal cholinergic interneurons,^35,36^ thereby modulating striatal output and, in turn, influencing pallidal activity. This pathway constitutes a key component of the mesocircuit proposed by Schiff.^10^ Previous studies have shown that unilateral high-frequency electrical stimulation of the CL can restore arousal and recruit fronto-parietal networks associated with consciousness.^17,37^ However, the gain-of-function experiments differ fundamentally from our pharmacological inactivation approach. Whereas electrical stimulation may activate local neurons together with fibres of passage and both orthodromic and antidromic pathways, muscimol selectively reduces local neuronal activity. These complementary approaches therefore probe different aspects of thalamic function and are not directly comparable.

Overall, these observations support a functional specialization within the intralaminar nuclei. Across both microstimulation and pharmacological inactivation, CM/Pf perturbation consistently produced stronger and more reproducible behavioural effects than comparable manipulations of the CL, providing causal evidence that these intralaminar nuclei are not functionally interchangeable. Rather, our findings suggest that the mesocircuit may rely on partially parallel functional pathways, including a more direct CL-thalamo-cortical pathway and a CM/Pf-striato-pallido-thalamo-cortical pathway involved in awareness. Within the mesocircuit framework, this dissociation may reflect complementary contributions of intralaminar nuclei to conscious access by modulating cortical networks either directly (CL) or indirectly through basal ganglia efferences (CM/Pf).

This observation is particularly important given that most clinical neuromodulation strategies have targeted the CL nucleus despite limited and heterogeneous outcomes, raising the possibility that insufficient engagement of thalamo-striatal pathways contributes to their inconsistent efficacy. Together, our findings highlight the heterogeneity of thalamic contributions within the mesocircuit and challenge the implicit assumption that all intralaminar nuclei contribute equivalently to consciousness-related processes.

Unlike lesion-based clinical studies, this preclinical NHP model enables causal and reversible manipulation of specific thalamic nuclei. Although subjective experience cannot be directly assessed, this approach provides a unique opportunity to investigate the respective contributions of intralaminar pathways to wakefulness and awareness-related processes within the mesocircuit framework.

### CM/Pf modulation engages frontal cortico-thalamo-striatal circuits

To investigate the large-scale networks associated with CM/Pf modulation, we combined local pharmacological perturbations with [¹⁸F]FDG PET imaging. This approach allowed us to characterize how focal CM/Pf modulation propagates across large-scale cortico-thalamo-striatal networks.

Bicuculline-induced activation of the CM/Pf produced marked and widespread increases in glucose metabolism within the left anterior striatum, including the putamen, caudate nucleus and ventral striatum, as well as bilaterally within the insula, orbitofrontal cortex, medial and limbic prefrontal cortices, and anterior cingulate cortex (ACC). Increased glucose metabolism was also observed bilaterally in the external pallidum and in the ventral posterior putamen. At the thalamic level, the strongest metabolic increase was observed at the injection site itself, namely the CM/Pf complex, but also extended to neighbouring nuclei including the MD, VPm, VA and VL. Importantly, VA and VL constitute major thalamic targets of basal ganglia output, supporting the interpretation that CM/Pf modulation recruits downstream pallido-thalamic components of the mesocircuit. In addition, bicuculline injections during PET acquisitions were associated with increases in heart rate and blood pressure. Although these physiological measures were not primary outcomes, they are consistent with previous evidence that CM/Pf activity influences autonomic and arousal-related networks alongside fronto-striatal mesocircuit pathway.

The widespread metabolic response is consistent with the known thalamostriatal projections of the CM/Pf^35,36^ and supports the view that this nucleus acts as a major hub linking basal ganglia output to distributed frontal cortical networks. The widespread fronto-striatal and thalamo-cortical metabolic changes observed following CM/Pf modulation are compatible with large-scale network interactions thought to support conscious access. Although [¹⁸F]FDG-PET does not directly measure conscious processing, the distributed pattern of cortical activation observed here is consistent with both the anterior correlate within the GNW model of ignition^38^ and the frontal connection of the mesocircuit.^9,11^ Given its extensive projections to associative and limbic territories^35^ and its major excitatory input to the striatum,^39^ the CM/Pf is anatomically well positioned to coordinate these distributed network effects. These findings are consistent with clinical observations in disorders of consciousness, where recovery is associated with restoration of fronto-striatal metabolism and connectivity. Taken together, these observations provide network-level evidence that CM/Pf activity influences multiple components of the mesocircuit through interconnected thalamo-striatal, pallidal and thalamo-cortical pathways.^40^

In contrast to bicuculline, muscimol injections, which suppress CM/Pf activity, induced hypometabolism in the thalamus, anterior striatal territories, and frontal cortical regions that showed increased metabolism following bicuculline administration. Similar hypometabolism in the thalamus has been reported in chronic DOC patients.^41^ Recent studies investigating the consequences of brainstem reticular formation injury have highlighted connectivity between the ACC and anterior insula as a potential cortical substrate contributing to the balance between arousal and awareness.^42^ Interestingly, these two cortical regions also exhibited opposite metabolic responses following bicuculline and muscimol injections into the CM/Pf. Although indirect, this observation is consistent with the hypothesis that ACC-insula interactions contribute to the regulation of arousal and awareness. More broadly, this experimental approach demonstrates how focal perturbation of a single intralaminar nucleus can reshape distributed mesocircuit activity at the whole-brain level. Together with the behavioural findings, these results support the hypothesis that CM/Pf contributes to mesocircuit dynamics through thalamo-striatal pathways associated with awareness, while wakefulness remained globally preserved.

### Limitations and perspectives

Despite the strengths of this study, several limitations should be acknowledged. First, the relatively modest behavioural effect may partly reflect the use of unilateral injections, which could allow compensatory mechanisms in the contralateral hemisphere. Further studies using bilateral muscimol injections may help determine the extent to which interhemispheric compensation influences these behavioural effects. Second, given the small size of the CL, whose thickness is less than 1 mm, we cannot exclude the possibility that muscimol diffusion extended into neighbouring thalamic nuclei, including the more medial MD or the more lateral VP, potentially attenuating or modifying the observed effects. Third, as in most non-human primate studies, the number of animals was necessarily limited. Accordingly, the present findings should be interpreted cautiously when extrapolating to patients with disorders of consciousness. Nevertheless, the NHP model provides a unique opportunity to causally investigate mesocircuit dynamics through selective and reversible manipulation of specific thalamic nuclei. The within-subject design further limits inter-individual variability and strengthens the robustness of the observed effects. These mechanistic insights complement previous clinical studies of thalamic deep brain stimulation in DOC,^23,24,43^ by helping refine the identification of candidate thalamic targets. Taken together, our findings provide a mechanistic rationale for extending current neuromodulation approaches beyond the CL and investigating the CM/Pf as a potential therapeutic target in DOC.

## Conclusion

In conclusion, this preclinical NHP study combining causal perturbations with [¹⁸F]FDG PET imaging identifies the CM/Pf as a major thalamo-striatal modulator of mesocircuit dynamics. Brain imaging further demonstrates how CM/Pf modulation influences widespread frontal, striatal, thalamic and pallidal components of the mesocircuit. Together, our behavioural and metabolic findings support the view that distinct intralaminar nuclei make complementary contributions to mesocircuit function, with the CM/Pf playing a particularly prominent role in thalamo-striatal mechanisms supporting awareness-related processes. These results refine current models of mesocircuit organization, provide a mechanistic framework for developing targeted neuromodulation strategies in disorders of consciousness, and support further investigation of the CM/Pf as a therapeutic target.

## Data availability

The data that support the findings of this study are available from the corresponding author, upon reasonable request.

## Acknowledgements

We thank Fidji Francioli, and Serge Pinede for animal care and technical assistance. In addition, we would like to thank Sophie Lancelot, and all members of CERMEP for MRI imaging.

## Supplementary material

Supplementary material is available at *Brain* online.

---

**Supplementary Figure 1:**
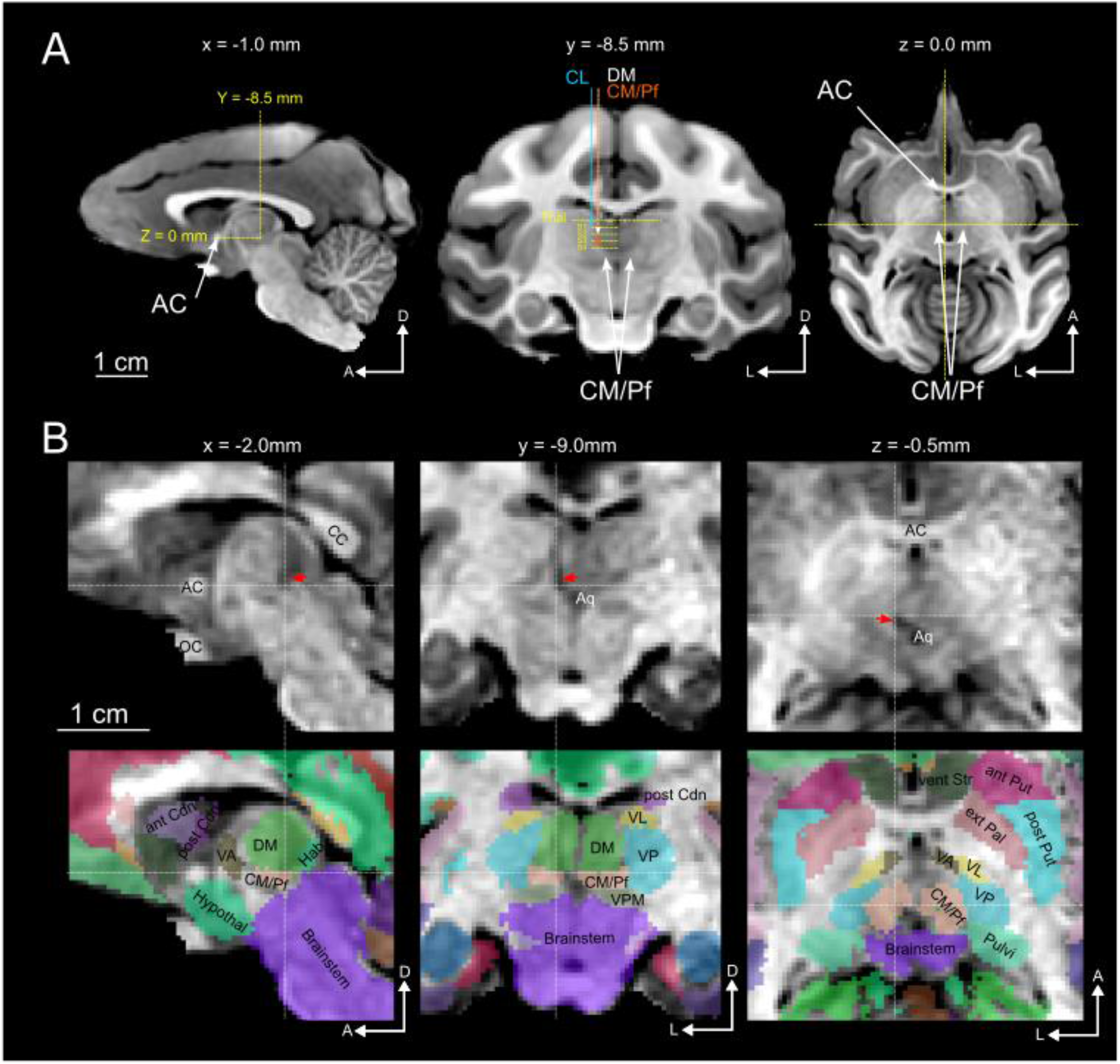
Anatomical localization of stimulation and injection sites. **(A)** Localisation of the injection and stimulation sites. The figure shows a T1weighted MRI image of MF2 recorded before surgery (3T scanner; CERMEP-Bron, France). The image is in the acpc native space where the origin is the posterior edge of the anterior commissure, and the image is rigidly rotated in order to ensure that horizontal plane is parallel to the anterior commissure-posterior commissure line (but distances were not normalized). The yellow dotted lines on the sagittal view represent the position of the frontal and horizontal slides. On the frontal view are represented the paths of the main electrode entrances (blue arrow for the CL stimulation sessions, white arrow for the MD stimulation sessions and orange arrow for the CM/Pf stimulation sessions. As the MD and the CM/Pf sessions used the same paths, the two arrows are on top of each other. The top yellow dotted line illustrates the depth of the thalamic entrance. S1, S2, S3, S4 are 4 depth positions at 1.5 mm of distance. (**B)** Illustration of a canula trajectory. The first row shows the average of two T1weighted MRI images from MF1, recorded during the first bicuculline PET session (in the acpc space, comparable to **A**). The trajectory is visible as a hyposignal trace passing through the left thalamus (red arrow). The dotted white lines illustrate the slice position of the views with different orientations. The slice coordinates are shown above the corresponding images. The second row shows the projection of the updated version of the Fascicularis atlas on the T1 weighted image. It shows the location of CM/Pf at the tip of the dark trace. *AC: anterior commissure; ant CdN: anterior Caudate Nucleus; ant Put: anterior Putamen; Aq: Aqueduct; CC: corpus callosum; CM/Pf: Centromedian-Parafascicular; ext Pallidum: external Pallidum; Hypothal: Hypothalamus; MD: Mediodorsal; int Pallidum: internal Pallidum; OC: optic chiasma; post CdN: posterior Caudate Nucleus; post Put: posterior Putamen; Pulvi: Pulvinar; vent Str: ventral Striatum; VA: Ventral Anterior; VL: Ventral Lateral; VP: Ventral Posterior; VPM: Ventral Posterior medial; Raphe: Raphe nucleus. The orientation directions are illustrated by the arrows (D= dorsal; A= anterior, L= left)*.

**Supplementary Figure 2:**
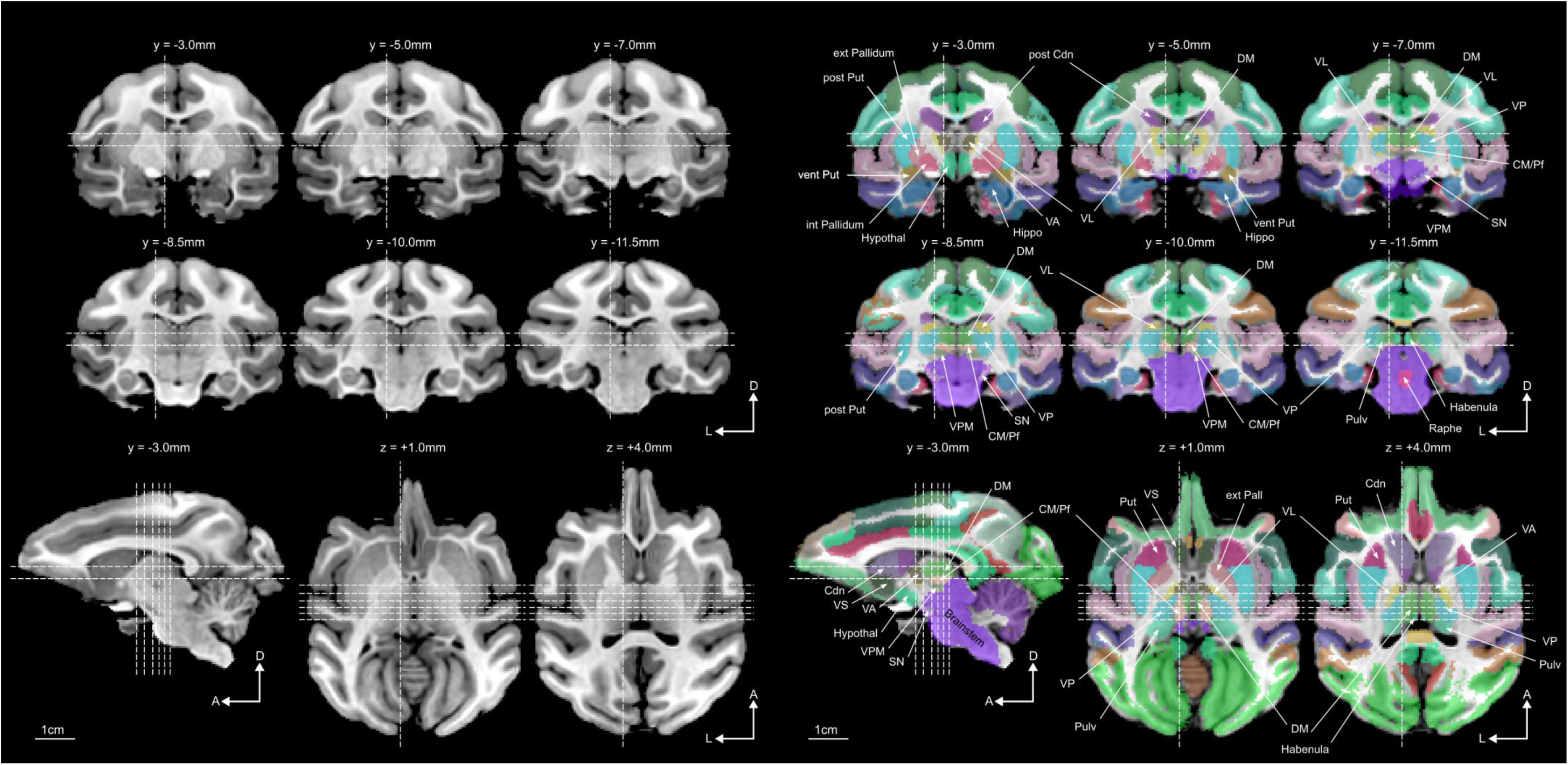
An updated version of the MaxProb atlas ^22^. The updated version of the maxprob atlas is projected on the same MRI image as Supplementary Fig. 2 (T1weighted MRI image of MF2 recorded before surgery with a 3T scanner). Only the modified structure has been labeled here. The dotted white lines represent the position of the views that have a different orientation. *ant CdN: anterior Caudate Nucleus; ant Put: anterior Putamen; CM/Pf: Centromedian-Parafascicular; ext Pallidum: external Pallidum; Hypothal: Hypothalamus; MD: Mediodorsal; int Pallidum: internal Pallidum; post CdN: posterior Caudate Nucleus; post Put: posterior Putamen; Pulvi: Pulvinar; vent Str: ventral Striatum; VA: Ventral Anterior; VL: Ventral Lateral; VP: Ventral Posterior; VPM: Ventral Posterior medial; Raphe: Raphe nucleus. The orientation directions are illustrated by the arrows (D= dorsal; A= anterior, L= left)*.

**Supplementary Figure 3:**
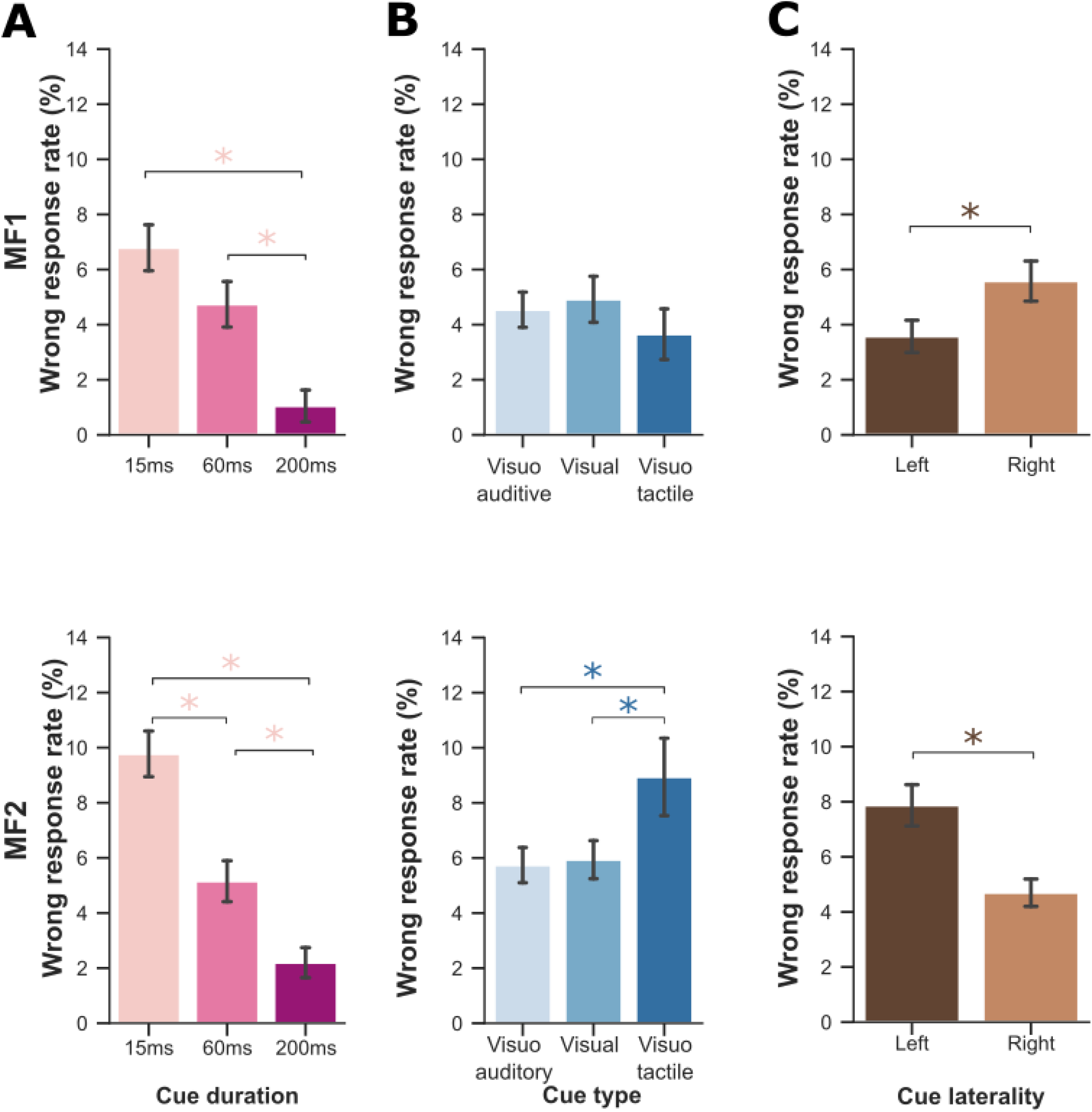
Influence of cue characteristics on wrong response rates during control sessions. Percentage of wrong responses (mean ± SEM) during baseline sessions as a function of: **(A)** cue duration (15, 60, or 200 ms); **(B)** cue modality (visual, visual-auditory, or visual-tactile); **(C)** cue laterality (left or right visual field). Shorter cue durations were associated with significantly higher wrong response rates (ANOVA, *F*(2,1812)=42.50), *P*<0.001), indicating increased task difficulty for brief stimuli. A significant main effect of cue modality was observed (*F*(2,1812)=3.03, *P*<0.05); however, post hoc comparisons did not reveal significant differences between individual cue modalities (all *P*>0.05). These findings indicate that behavioural performance was primarily influenced by cue duration rather than sensory modality.

