## Supplementary Material and Methods for "CM/Pf contribution to mesocircuit dynamics underlying awareness in non-human primates"

#### Surgery procedure

By positioning the center of the chamber on the midline and opening a bony window of 6 mm on either side of this median line and 4 mm in front and posterior of this center, the area of interest became accessible for the 3 thalamic nuclei. The CL, the narrowest structure (<1mm thick) is 4 mm from the midline, while the penetrations for the MD and the CM/PF complex were made at lateral 2 and 3 mm from the midline. After the implantation surgery, a second imaging session was performed to really locate the area of interest. Before starting the real experimental period starting with the stimulations, we made a series of electrophysiological recordings to determine the coordinates of the thalamic entrance (after the passage of the ventricles) and to determine the depth of our stimulation and injection sites: for the CL and the MD, the stimulations were done at three dephts (at 1.5 and 3 and 4.5 mm after the thalamus input) while for the CM/PF located at 6 mm. In the CM/PF, stimulation was done at 2 laterality and 2 anteriority, the same than both MD and CL.

#### Behavioural task

Presentation (Neurobehavioral systems, Inc, MA, USA) and Scenario Manager (Institut des Sciences Cognitives Marc Jeannerod, CNRS, Bron, France) software were used to control the behavioral task, along with solenoid valves that opened and closed the reward delivery and the air-puff system.

An optic-fiber system was used to detect the presence of the hand on a lever in front of the monkey's chair. The animal was seated in a closed primate chair, only open at left hand allowing the animal access to touch the screen and perform the task (Figure 1a). The monkey must keep its hand on the lever until the cue is displayed, and then until a target is displayed. The monkey can then remove his hand from the lever and touch the target. If the monkey selects the target corresponding to the cue, reward was delivered (0,36mL/trial for MF1, 0.47mL/trial for MF2). If the monkey selects the target on the other side, it gets no reward. If he presses elsewhere on the screen, does not raise his hand from the bar or raises his hand from the bar before the targets appear, these are considered errors, and the animals gets no rewards. An early release corresponded to the premature removal of the hand from the lever, and a non-target touch was defined as a touch on the screen outside the target areas.

In the visual condition, the cue was represented by a gray square, which was intentionally difficult to detect. The auditive conditions is a gray square with a sound emanating from two loudspeakers that were placed on either side of the animal. The sound has a frequency of 450 Hz. At last, the somaesthetic condition is a breath of air, called a puff, fed through two plastic tubes and placed 5 cm from each of the monkey's cheeks. The puff was at 0.5 bar of pressure, controlled by a control valve. In the first two conditions (Figure 1a), three times of cue were used: 15ms, 60ms and 200ms. For the tactile condition, only a duration of 15ms were used, because the puff can be aversive for when delivered with excessive pressure or duration. This task was consisting in 4 blocks of 112 trials. In each bloc, we have 8 trials of each sub-conditions and on each side of the screen (right or left), so 16 trials of each sub-conditions. So, in total, there are 14 possibilities: 3 delays on 2 different sides for the visual, 3 delays on 2 different sides for the audio, and 1 delay on 2 different sides for the puff.

### Microstimulation

For the MD and CL territories, two anterior positions (AC -8 to -9; 1 mm apart) were tested on the same laterality, each at three different depths spaced 1.5 mm apart with the first stimulation site, 1.5 mm inside the thalamus by entering through the lateral ventricles (Fig S1). This resulted in a total of six stimulation sites per territory and animal. Each site was stimulated twice. For the CM/PF territory, the same two anterior positions used for the MD and CL and two lateralities were tested (Lat 2 and 3 for the Pf and CM nucleus, all at the same depth (around 6 mm below the thalamic entrance). The CM/PF depth was checked by neuronal recording to be sure to stimulate the center of this small structure (1.5 to 2 mm thick). This resulted in four stimulation sites per animal, with two stimulation sessions per site.

Microstimulation were a For the tactile condition, onlpplied during the task, at 200 µA, in bipolar configuration, frequency 130 Hz and amplitude 1V. The same tungsten microelectrodes used for neuronal recording was used for the microstimulation (Phymep microelectrode, impedance: 0.7–1.5 MOhm measured at 1000 Hz). Stimulation takes place for 600 ms before the cue. At the end of the task, animals underwent a 6-minute period comprising three repetitions of 1-minute stimulation alternating with 1-minute rest intervals.

### Microinjections

Microinjection sessions were performed once daily, twice per week, with at least one control day between sessions, to allow for a wash-out period and to minimize the animal's discomfort. For the CM/PF territory, the same two anterior positions used for the stimulation were tested on the same side and at the same depth for each animal, resulting in two injection sites. Each site received two injections. For the CL territory, protocols differed between animals. In MF1, two anterior positions were tested on the same side and at the same depth, corresponding to two injection sites, with two injections per site. In MF2, two anterior positions were tested on the same side; however, one position was tested at two different depths (1.5 mm apart), resulting in three injection sites. One site received two injections, while the other two sites each received a single injection.

#### Eye-tracking

Eye state (open vs. closed) was continuously recorded using an infrared eye-tracking system (ETL-200, ISCAN Inc., Woburn, MA, USA) sampling at 240 Hz. The vertical eyelid aperture was extracted offline and binarized using a predefined amplitude threshold to classify each sample as eyes open or eyes closed. From this binary time series, the probability of eye closure was computed as the proportion of samples classified as "closed" within a given time window. This measure was analysed across experimental days, yielding time-resolved and day-resolved estimates of eye-closure probability.

#### PET imaging analysis

PET imaging was performed using a 3T Siemens Biograph mMR simultaneous PET-MRI scanner. Attenuation was obtained using HiRes MRI sequence. PET emission images were reconstructed using the PSF (Point Spread Function) algorithm with 8 iterations, 21 subsets and a zoom factor of 3, over 41 dynamic frames (4×15s, 4×30s, 6×60s, 27×180s). Reconstructed volumes were 127 slices (2.032 mm thickness, 256 x 256 matrices of 0.9 x 0.9 $mm^2$ voxels). PET scans followed the same procedure for each individual. Rigid realignment between the frames to the average PET image without considering the first 7 frames because of weak signal. After realignment, new mean PET images were registered to their corresponding individual anatomical MRI, which was registered to the *macaca fascicularis* MRI template [1] . A new version was performed to include thalamic nuclei (Deroche et al, 2020). The subdivision of the thalamus into six nuclei (VA,VL and CM/Pf complex, MD, VP and Pulvinar) was based on

Lanciego and Vazquez's histological atlas [2]. Figure S2 presented manual refinements of the thalamic nuclei, hypothalamus and raphe nucleus). The thalamic nuclei boundaries were identified based on three studies [3,4,2] with the help of 3 different macaque atlas projected on the T1w images : the 2020 version of the maxprob atlas, the SARM atlas [5] and the D99 atlas [6]. The boundaries of hypothalamus were manually modified after projecting the SARM atlas on the individual data [5]. The present hypothalamus region corresponds to the tuberal hypothalamus and the posterior hypothalamus (SARM atlas, third level of segmentation). The same methodology was used for building the maxprob atlas. The 4 individual atlases were normalised on the macaca fascicularis template. and fused. Only the voxels in which the data from the four individus converged (all 4 pointing to the same label) were preserved and added to the 2020 version of the maxprob atlas (Deroche et al, 2020). Dice index was used to evaluate the accuracy of automated segmentation with the new atlas versus manual segmentation [1].

The Raphe region was functionally identified in a previous PET study using the [$^{18}$F]MPPF tracer of the serotonin system and the Macaca Fascicularis template [7]. The Raphe region added to the maxprob atlas corresponds to the inclusive mask created from the 70% upper signal within the brainstem in non-displaceable binding potential ($BP_{ND}$) parametric PET image.

Transformations from native PET to individual MRI and individual MRI to template were then concatenated to provide direct (and inverse) affine and nonlinear transformations from PET native spaces to the template space. Each voxel value was multiplied by the injected tracer dose (in MBq) and divided by the individual's weight (in g) to compute SUV (standardized uptake value). The frames were then averaged per block of 15 minutes to end up with 6 images per session. The results were obtained from the last block of parametric PET images (between 75 min and 90 min after injection). The images were skull stripped using an inclusive brain mask and each voxel value was divided by the mean value of the brain. The parametric images were then transformed to the common template space with linear interpolation. Region delineation was achieved with an updated version of the *macaca fascicularis* maximum probability atlas [1] back projected on the parametric images in the native space. Regional parametric values were obtained by calculating the mean per label of the atlas.
